# *PathogenSurveillance*: an automated pipeline for population genomic analyses and pathogen identification

**DOI:** 10.1101/2025.10.31.685798

**Authors:** Zachary S. L. Foster, Martha A. Sudermann, Camilo Parada-Rojas, Logan K. Blair, Fernanda Iruegas-Bocardo, Upasana Dhakal, Ricardo I. Alcalá-Briseño, Hung Phan, Tori R. Schummer, Alexandra J. Weisberg, Jeff H. Chang, Niklaus J. Grünwald

## Abstract

Whole genome sequencing (WGS) offers a comprehensive, organism-agnostic method that effectively meets the need for efficient, reliable, and standardized responses to emerging threats from pathogens and pests. Here, we present *PathogenSurveillance*, an open-source and automated Nextflow pipeline for population genomic analyses of WGS data. It is designed with features tailored for biosurveillance and is suitable for in-field or point-of-care diagnostics. *PathogenSurveillance* is flexible, accommodating short- and long-read datasets and mixed samples of prokaryotes and/or eukaryotes. It automates all steps, including reference identification and retrieval from the NCBI Assembly database, and produces customizable interactive reports with summaries, phylogenetic trees, and minimum spanning networks that enable species and subspecies level identification. It also outputs quality control metrics and carefully names and organizes output files to facilitate downstream analyses. The pipeline runs on any Linux-based system and minimizes the need for advanced computational expertise. Source code is available on GitHub under the open-source MIT license. The pipeline expands the toolkit for real-time biosurveillance, enabling rapid detection and monitoring of pathogens and pests for rapid response to novel variants.

## Introduction

Emerging invasive pathogens and pests pose an escalating global threat, largely driven by anthropogenic factors such as intensified international trade (Meyerson and Mooney 2007; Hulme 2009; Fisher et al. 2012; Liebhold et al. 2012). Their sudden appearance and potential to spread rapidly can outpace existing biosurveillance systems, making it difficult to mount timely responses (Carvajal-Yepes et al. 2019). Consequently, diseases caused by these novel organisms may reach epidemic levels before effective diagnostic methods are developed. To address this challenge, biosurveillance efforts require faster and more reliable detection that enables timely interventions (Gardy and Loman 2018; Hamelin et al. 2022). This is especially relevant for OneHealth frameworks, which emphasize the interconnected movement of pathogens across human, animal, and environmental domains, as seen in cases of antimicrobial resistance (Djordjevic et al. 2024).

Whole Genome Sequencing (WGS) is an important tool for biosurveillance, particularly in the early detection of infectious disease outbreaks (Weisberg et al. 2021). Methods for analyzing WGS data are highly effective for detecting emerging or reemerging invasive pathogens and pests, as they do not require *a priori* knowledge or optimization for specific organisms and can be applied to diagnosing rapidly developing threats even when the causative agent is unknown (Gardy and Loman 2018; Weisberg et al. 2021). However, key hurdles to the adoption of WGS are the need for appropriate infrastructure and domain expertise in both computational methods and pathogen biology (Oakeson et al. 2017; Iles et al. 2021). Thus, it is essential to design computational workflows that directly confront these challenges and empower users without specialized training to work with WGS data for disease detection.

WGS-based approaches commonly used for characterizing genetic clustering can be broadly grouped as *k*-mer, multigene, or variant-based phylogenies. *K*-mer based methods have the advantage of being extremely fast and computationally efficient (Pierce et al. 2019). They work by comparing the presence of only exact sequences of a given length (known as *k*-mers). The *k*- mer contents of genome sequences can be summarized by a systematically selected subset of hashed *k*-mers called a sketch and compared to a database of other sketches corresponding to reference genomes (Tian et al. 2020; Irber et al. 2022; Irber et al. 2024). However, sketch-based approaches are limited in their ability to infer taxonomic breadth, as their reliability declines with decreasing genome sequence similarity (Ondov et al. 2016). Conversely, multigene phylogenies can be used to group samples at various taxonomic levels, making such methods ideal for placing organisms of unknown identities in a phylogenetic context. But these approaches have limited resolution, are less scalable, and require domain knowledge to select appropriate marker gene and reference sequences. The major strength of variant-based (single nucleotide polymorphisms (SNPs)) analyses lies in their high resolution, which enables clustering of closely related samples to identify clonal relationships and reveal potential transmission events. Variant-based approaches have similar limitations as multigene phylogeny-based approaches and are even more sensitive to reference selection and assembly quality (Weisberg et al. 2021).

Various pipelines have been developed for pathogen surveillance. *GATK PathSeq* enables identification of pathogens from host-derived samples and is highly efficient, integrating well with standard computational environments, although it requires a reference database of potential microbes and host sequences (Walker et al. 2018). *Bactopia* and *Galaxy@Sciensano* are collections of seamlessly integrated tools designed for analyses of prokaryotes, but they require domain knowledge to select appropriate reference sequences (Petit and Read 2020a; Bogaerts et al. 2025). *CZ ID* (formerly *IDseq*) is a web-based pipeline for analyzing metagenomic data for pathogen detection but does not provide tools for multigene phylogeny construction or variant analyses (Kalantar et al. 2020). *Pathogenwatch* is a web service that supports phylogenetic analyses, primarily through Multi-Locus Sequence Alignment (MLSA) based phylogenetic approaches, and has utility for in-depth analyses (Argimón, David, et al. 2021; Argimón, Yeats, et al. 2021; Sánchez-Busó et al. 2021). However, *Pathogenwatch* is not optimized for use on local machines or private clouds and is largely limited to specific pathogen groups. *PathoGFAIR* provides a Galaxy-based workflow to detect and track pathogens from metagenomic Nanopore sequencing (Nasr et al. 2025).

Here we present nf-core/*PathogenSurveillance,* an open-source pipeline for automated population genomic and evolutionary analysis. This pipeline was released within nf-core, a collaborative, community-driven framework for developing and sharing standardized pipelines built on the Nextflow workflow management system (Di Tommaso et al. 2017). *PathogenSurveillance* leverages nf-core modules and community peer review to ensure robust functionality, while its implementation in Nextflow provides scalability, reproducibility, and portability. *PathogenSurveillance* automatically processes and analyzes short- and long-read WGS data along with publicly available genome assemblies. It features specialized capabilities for pathogen and pest identification, including flexible support for user-defined or automated reference sequence identification and retrieval, as well as the ability to concurrently analyze mixed prokaryotic and eukaryotic samples and populations. Other distinctive features include the integration of *k*-mer-, multigene phylogeny-, and variant-based approaches to deliver multiple levels of phylogenetic resolution for pathogen identification. The interactive output presents easily interpretable results, facilitating actionable responses. Moreover, output files are organized for deeper downstream investigation, extending use of *PathogenSurveillance* to population genomic studies to a wider range of organisms. To validate and illustrate its utility, we applied *PathogenSurveillance* to test datasets, demonstrating accuracy in automated reference selection and sample identification.

### Implementation

We developed *PathogenSurveillance* for WGS data analyses of pathogens or pests. This pipeline is user-friendly and automates all steps from reference sequence selection and retrieval to the generation of interactive reports detailing sample identity, phylogenetic relationships, and population structure (Figure 1). The pipeline was designed according to widely accepted core principles, with a particular focus on minimizing the need for advanced computational expertise by end users and incorporating specialized features for pathogen surveillance. *PathogenSurveillance* follows the FAIR (Findable, Accessible, Interoperable and Reusable) principles for processing data using workflows (De Visser et al. 2023).

**Figure 1.**
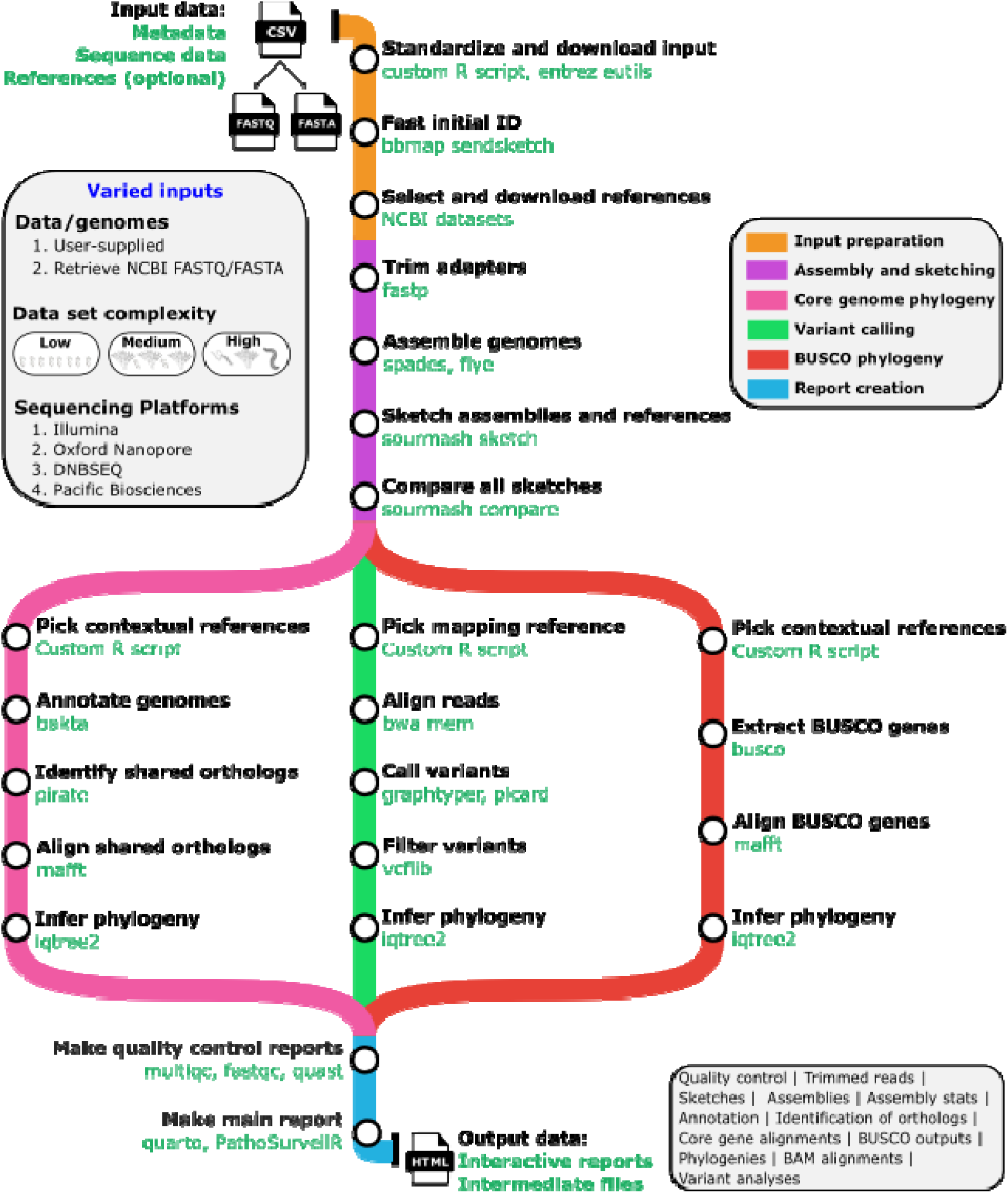
*PathogenSurveillance* automates comprehensive analysis of prokaryotic and eukaryotic whole genome sequencing data. A TSV file indicating paths or URLs to FASTQ or FASTA files of short or long read sequencing data initiates the process of downloading and formatting of input files and references. The next part of the main branch quality controls sequencing reads and generates assemblies as well as *k-*mer sketches. It then diverges into three branches. The left branch is for samples of prokaryotic origin and will generate core phylogenies based on all shared orthologous gene sequences and variants, respectively, for each group of related samples. The right branch is for samples of eukaryotic origin and will build a phylogenetic tree for each group of related samples based on shared, single-copy BUSCO gene sequences. The middle branch creates SNP phylogenies for both prokaryotic and eukaryotic samples. Finally, the branches converge and *PathogenSurveillance* will generate a final HTML report with interactive plots and tables as well as quality control reports. Prominent steps in the workflow are printed in black beside each node with select corresponding tools used printed below in green.

*PathogenSurveillance* streamlines installation by leveraging Nextflow, an open-source software tool and domain-specific language that simplifies the creation and execution of data-intensive, scientific computational workflows (Di Tommaso et al. 2017). Nextflow uses container frameworks, such as Docker, Apptainer, or Conda, to automatically and reproducibly install all required dependencies as needed upon first execution. Each process runs in its own isolated container and file system, eliminating software incompatibilities and ensuring consistent behavior across any Linux-based environment. As part of *PathogenSurveillance*’s development, we contributed seven new modules and made updates or bug fixes in eight existing nf-core modules (Supplementary Table 1).

The input process is also streamlined without sacrificing flexibility or configurability. *PathogenSurveillance* minimally requires a tab-separated value (TSV) file as input, with the only mandatory field being source of sequence data. This field can be populated with any combination of paths to local files or URLs to remote files, NCBI Short Read Archive (SRA) accession numbers, or NCBI Biosample IDs associated with SRA accessions. Input data may correspond to a single sample, multiple taxonomically disparate samples (including both prokaryotic and eukaryotic organisms), populations of related samples, or any combination thereof. While *PathogenSurveillance* also has flexibility in analyzing samples with raw reads derived from Illumina, PacBio, or Oxford Nanopore technologies, it does not support data combined for the same sample from multiple sequencing platforms. For the latter case, users will need to select one source of read data. All other metadata for samples are optional but recommended if available, to support downstream analyses. Users can also provide a second optional TSV file to specify reference genomes. Furthermore, users can indicate whether user-defined reference genomes should be included along with automatically selected references or used as the sole source, thereby excluding other possible references from analyses.

The pipeline is initiated with a single command. Upon initiation, it automatically generates a *k-*mer sketch from each sample’s raw reads and compares these to sketches of all of NCBI RefSeq using bbmap sendsketch (Figure 1) (Bushnell 2014). If chosen by the user, it then identifies and retrieves assemblies that are possible reference sequences. Next, *PathogenSurveillance* processes and assembles sample reads and generates new *k-*mer sketches using sourmash to infer Average Nucleotide Identity (ANI) values that can be used to refine reference selection (Konstantinidis and Tiedje 2005; Pierce et al. 2019). The pipeline selects the next workflow branches based on whether the sample is inferred to be of prokaryotic or eukaryotic origin. For prokaryotic datasets, it will annotate genome assemblies, generate ortholog clusters, identify shared orthologs among samples and their references, and construct a core genome phylogenetic tree. If the pipeline identifies clusters of closely related samples, it will map reads to their common reference assembly, call SNPs, and generate a SNP tree as well as a minimum spanning network. For eukaryotic datasets, *PathogenSurveillance* does not annotate genome sequences, as automated annotation is impractical because taxonomic groups require specialized training datasets. Instead, the pipeline generates a phylogenetic tree based on identified Benchmarking Universal Single-copy Orthologs (BUSCO) sequences (Manni et al. 2021). A final interactive report is generated as an HTML file wherein users can define groups and adjust visualizations. All intermediate files, such as assemblies and annotations, are stored and made accessible for additional analyses (Supplementary Figure 1). *PathogenSurveillance* also produces quality control reports for reads and assemblies. Details on key workflow steps of the pipeline are provided in the following sections. The report created lists all the references and sources for computational tools used in the analysis (Sample reports can be downloaded at https://github.com/grunwaldlab/pathogensurveillance_publications/tree/main/publication/example_reports).

*PathogenSurveillance* is portable and can be executed on any Linux platform, including local desktops, high performance computing (HPC) clusters, and commercial cloud environments. In addition, *PathogenSurveillance* can be launched from Sequera Platform, a web-based platform. A comprehensive guide to executing the pipeline, including input formats and optional parameters, is available online (https://nf-co.re/PathogenSurveillance).

### Automated reference selection and population-level analyses

Effective selection of reference genome sequences is critical for accurate pathogen identification (Grünwald and Goss 2011; Weisberg et al. 2021). However, the likely taxonomic classification of a sample may not be readily known, particularly for emerging pathogens and pests. To mitigate this limitation, we implemented a rule-based system for context-aware selection and retrieval of reference genome sequences that balances per-sample accuracy with the potential for coherent population-level analyses (Figure 2). By default, *PathogenSurveillance* attempts to automatically detect and group related samples within the analyzed dataset. Users also have the option of manually assigning references to samples or specifying which subgroups of samples should be analyzed together.

**Figure 2.**
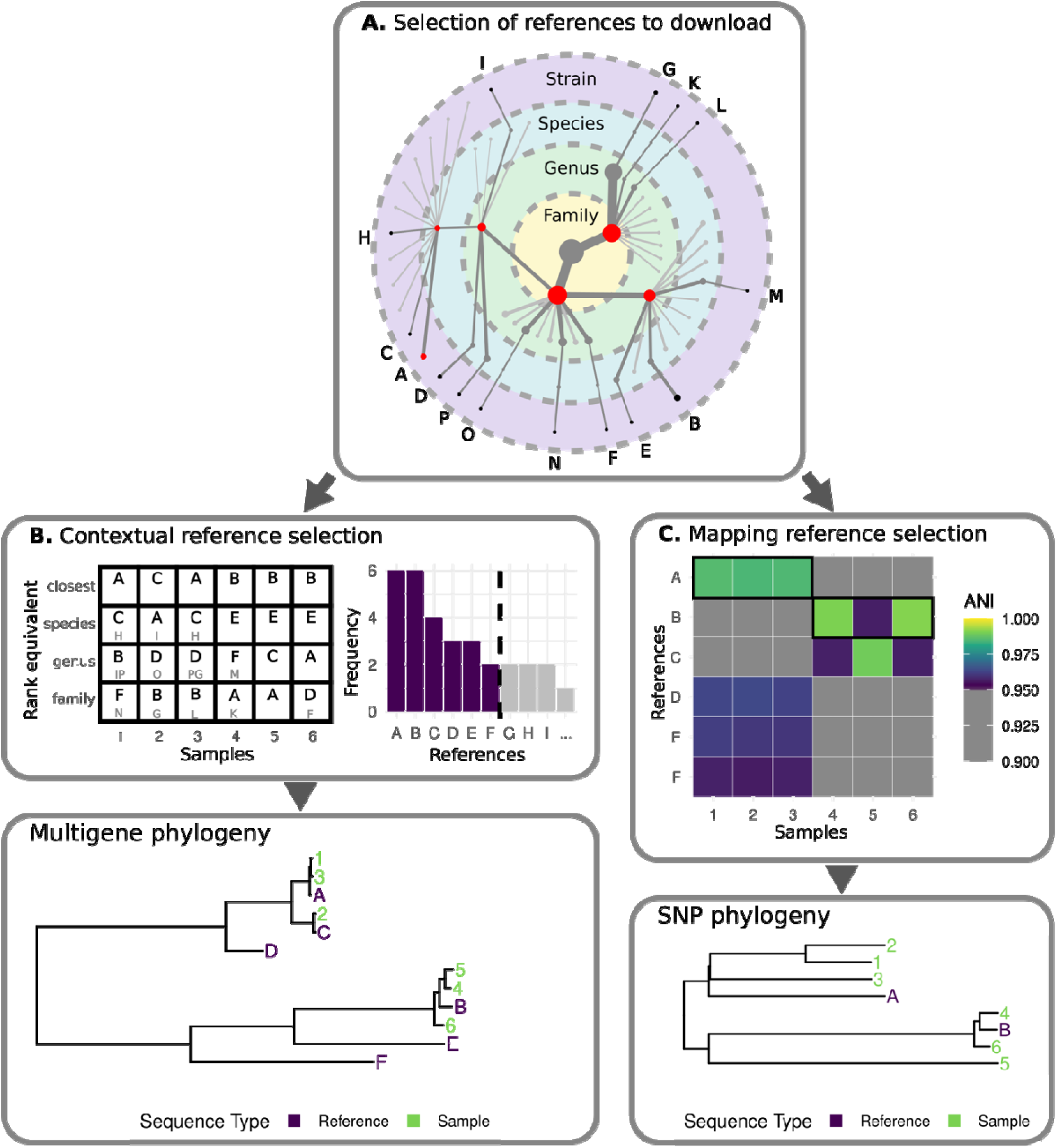
*PathogenSurveillance* automates reference selection for phylogenetic analyses. Shown is a hypothetical example illustrating sampling of genomes from **(A)** publicly available references for **(B)** multigene phylogenies and **(C)** SNP-based phylogenies or minimum spanning networks, respectively. **(A)** First, *PathogenSurveillance* compares *k-*mer sketches of raw reads to sketches of all NCBI RefSeq assemblies using BBmap sendsketch. Taxa likely present in samples are identified based on match similarity metrics (shown as red nodes in the taxonomic tree in (**A**)) and representative assemblies of subtaxa are downloaded for each detected taxon (black nodes). The pipeline prioritizes enrichment of candidate references from more closely related taxa but also includes more distant taxa in case the initial identification was incorrect. **(B)** From the set of references downloaded a subset are selected to provide context for multi-gene phylogenetic trees based on pairwise ANI estimates between samples and references. References estimated to be most similar to each sample are always included in addition to a minimal set of references selected for the set of samples as a whole that occur at a range of similarities to each sample. Each cell in the matrix represents a role that references can fit into. References are ordered by how many cells each can fill, as is shown in the histogram, and selected in this order until all cells are filled. **(C)** Mapping references are also selected for SNP calling based on estimated ANI values. A conceptual heatmap illustrates pairwise ANI comparisons, with cells exceeding the minimum threshold required shown in color. The reference with an estimated ANI exceeding a threshold for the most samples is chosen and this process is repeated until all samples have an assigned reference. Ties are broken based on mean ANI. A conceptual SNP-based phylogenetic tree demonstrates that only clusters with mapping references A and B met the minimum group size criteria for SNP calling and phylogenetic inference.

As described in the overview, rules are executed in two stages (Figure 1). In the first stage, a subset of possible reference assemblies relevant to the aggregated set of samples are downloaded (Figure 2A). For every unique family identified for samples analyzed, *PathogenSurveillance* will download the corresponding NCBI Assembly metadata, used to further refine reference selection based on genera, species, and strain information. If multiple references are available for a selected taxon, selection is additionally based on the prevalence of the associated taxon in the database indicating it is typical by NCBI standards, whether the assembly corresponds to a type strain, is in RefSeq, and associated with a Latin binomial. Additional criteria include assembly features such as assembly level, predicted assembly coverage of the genome sequence, absence of contaminating reads, and contig L50. In the second stage, the pipeline uses estimated ANI values to identify two types of reference assemblies among those downloaded (Figure 2). Contextual references are of well-characterized organisms selected to represent different taxonomic levels for phylogenetic analyses while mapping references are used for SNP calling.

To facilitate population-level analysis, the pipeline identifies contextual references that are shared across samples. This reduces the total number of distinct references and enables grouping of related samples for phylogenetic inference (Figure 2B). References are first partitioned into equal-sized bins based on a rescaled estimated ANI value for each sample. References most similar to each sample and those specified by the user are selected as initial candidates. *PathogenSurveillance* then iteratively selects additional references that best serve the entire group of samples. In each iteration, it prioritizes references that span the greatest number of bins across all samples, recalculating bin coverage after each selection. This process continues until every bin is filled by at least one selected reference. This process also helps reduce runtime and computational resources needed to construct the phylogeny.

For prokaryotic samples, core gene sequences are used to construct a phylogeny while for eukaryotic samples, BUSCO gene sequences are used (Manni et al. 2021). If *PathogenSurveillance* determines that too few gene homologs are shared among samples, it partitions the dataset into the fewest possible number of clusters such that each cluster meets the minimum threshold of shared gene homologs required to support robust phylogenetic inference. The default threshold is set to 10 gene homologs.

Mapping reference assembly quality and phylogenetic distance from samples affects the accuracy of SNP calling after read mapping (Iruegas-Bocardo et al. 2023). Moreover, only samples whose reads are aligned to the same mapping reference assembly are directly comparable in SNP analyses. *PathogenSurveillance* addresses these limitations by using estimated ANI values to cluster samples with their most closely related references in a sequential manner (Figure 2C). The process begins by selecting a reference that exceeds a defined minimum similarity threshold and is shared by the largest number of samples. When multiple references meet estimated ANI thresholds for the same number of samples, the one with the highest average similarity across those samples is chosen. This process is repeated iteratively until no samples can be matched to a reference. For each resulting cluster containing two or more samples assigned to the same reference, a pangenome graph method is executed to call SNPs (Eggertsson et al. 2019). This method is more robust to variation in phylogenetic distance and reference sequence quality (Eggertsson et al. 2019; Weisberg et al. 2021; Iruegas-Bocardo et al. 2023). SNP data are then used to create phylogenetic trees or minimum spanning networks.

### *PathogenSurveillance* outputs

*PathogenSurveillance* produces a rich set of data that is synthesized in tables and plots in interactive HTML reports for each user-defined group of samples (See for example: https://github.com/grunwaldlab/pathogensurveillance_publications/tree/main/publication/example_reports). A sunburst plot provides a summary of taxonomic classifications of the samples, predicted based on initial *k-*mer sketches of raw reads and is associated with a table that presents the statistics for each match. These preliminary classifications, based on analysis of raw reads, serve to guide *PathogenSurveillance* in the next round for selecting reference assemblies. Next, estimated ANI and Percent of Conserved Proteins (POCP) values are displayed as both interactive heatmaps and in a table of best matches for each sample (Qin et al. 2014). These give essential context for interpretation of predicted taxonomic identities, as ANI reflects similarity across homologous regions while POCP captures overall genomic similarity between compared genome sequences (Weisberg et al. 2021). Multilocus, phylogenetic trees provide taxonomic placements of samples (Minh et al. 2020). A minimum spanning network constructed from genetic distances is used to visualize clonal lineages within populations (Kamvar et al. 2014). Tables and plots are accompanied by expandable tooltips that explain how to interpret data and outline methods used.

In addition to the main report, *PathogenSurveillance* writes results from each step into structured folders that can be mined further for downstream analyses (Supplementary Figure 1). These output folders are organized hierarchically and named intuitively. Likewise, output files corresponding to specific samples, references, or groups of samples and references are consistently named based on user-definable identifiers. The organization and naming are intentionally designed to be both human-readable and machine-parseable, facilitating seamless integration into downstream analyses. The kind of analyses used by *PathogenSurveillance* necessarily output large and numerous files, not all of which are equally likely to be useful for downstream analysis. Nearly all intermediate files that take up little space are included in the output, but care has been taken to only include large files if their potential utility justifies their storage footprint. For example, quality filtering of BAM and VCF files occurs in several steps, each of which produces large files, but only the raw and final filtered versions of BAM and VCF files are preserved in the output. Additionally, by default all files in the output directory are links to files stored in a cache rather than the files themselves. This is the same cache that is used to resume pipeline runs and skip redundant steps. This has the benefit of only storing a single file when multiple runs of the pipeline produce the same file, as is common when testing multiple variations of a given dataset or pipeline parameters. Together, the careful curation of output files combined with the deliberately streamlined design significantly reduces the need for disk space, enabling larger analyses and efficient reanalysis of changing datasets.

The outputs also include quality control reports that support the interpretation of results. *PathogenSurveillance* automatically executes several standard quality control programs. FastQC is used to report basic and per read statistics for short read datasets while NanoPlot serves the same purpose for long read datasets (De Coster and Rademakers 2023). The adapter-trimming software, fastp, provides detailed reports on read filtering, trimming, and overall read quality (Chen et al. 2018). Quast is used to assess assembly quality metrics such as contiguity and completeness (Gurevich et al. 2013). BUSCO provides an additional estimate of genome assembly completeness by reporting the presence/absence of expected core genes. Finally, MultiQC is used to compile results from each of these programs into a single, comprehensive report, presenting quality control data for all samples in a consistent and well-organized format (Ewels et al. 2016).

### Additional user-friendly features

*PathogenSurveillance* incorporates extensive parallelization, error recovery mechanisms, and caching of intermediate results, enabling improved efficiency and resilience in automated execution of large-scale, long-running analyses. Parallel processing occurs as system resources and API limits allow. In HPC clusters and commercial clouds, this can scale to hundreds or thousands of jobs running in parallel, each using all available central processing units (CPUs) on its assigned node or virtual machine. Processes that encounter errors are automatically retried a set number of times, with each attempt incrementally increasing Random Access Memory (RAM) and CPU allocation to accommodate resource demands. Processes that communicate with external web services will pause before retrying after a failed attempt, allowing time for temporary network issues to resolve. Information about pipeline execution, including any warnings or errors encountered during the process, are presented in output reports.

Caching of intermediate results by *PathogenSurveillance* offers three key advantages. First, if the pipeline terminates prematurely either due to an excessive number of failed processes or a user-issued command, it can be restarted from the point at which it stopped. This is made possible by using the "-resume" option. Second, caching gives users opportunities to incorporate additional data or adjust parameters. Only the affected portions of the analysis are recomputed and as a result, rerun executions are dramatically faster. Third, and similarly, caching enables users to iteratively refine evolutionary and population-level analyses. Users can first run *PathogenSurveillance* without specifying references, relying exclusively on automated reference selection, and then manually choose more appropriate references after inspecting results. Reruns will be executed quickly and potentially be more robust. A drawback of caching is its large storage footprint, as seen in other comprehensive pipelines such as *Bactopia* (Petit and Read 2020). We mitigated this to some extent by designing the pipeline to intentionally avoid storing redundant intermediate files.

### Validation of *PathogenSurveillance*

We used various datasets to evaluate the accuracy and performance of *PathogenSurveilance.* First, a subset of publicly available *Serratia* genome sequences were used to test the accuracy of automated phylogenetic inference (Williams et al. 2022). In the original work, authors used two methods to quality control sequence data and used two different methods to assemble long-read PacBio and short-read Illumina data. From their original set of 664 samples, we manually selected a subset of 302 that captured the species and ecological diversity of the full set (Supplementary Table 2). The subset represented 11 species identified previously, with sequencing reads generated using long- and short-read technologies. We input their accession numbers and metadata into a TSV file. We also manually selected accessions of reference assemblies that were used in the original study (Williams et al. 2022). The pipeline was executed with a single command and in a mode that prevents *PathogenSurveillance* from automatically identifying and retrieving additional reference assemblies:

~~~
nextflow run nf-core/*PathogenSurveillance* -r 1.1.0 -resume - profile docker,serratia_full --outdir serratia_results
~~~

*PathogenSurveillance* generated a phylogenetic tree, along with all the other associated intermediate assemblies, annotations, and ortholog cluster files as well as quality control metrics.

The resulting tree was concordant in topology to the core gene phylogeny previously reported, and importantly, the species and lineage assignments were highly similar between the two analyses, with minor discrepancies restricted to placements of samples within clonal lineages (Figure 3) (Williams et al. 2022). Overall, evaluation of this test dataset demonstrates that with minimal user input, *PathogenSurveillance* can generate reliable phylogenetic and taxonomic insights.

**Figure 3.**
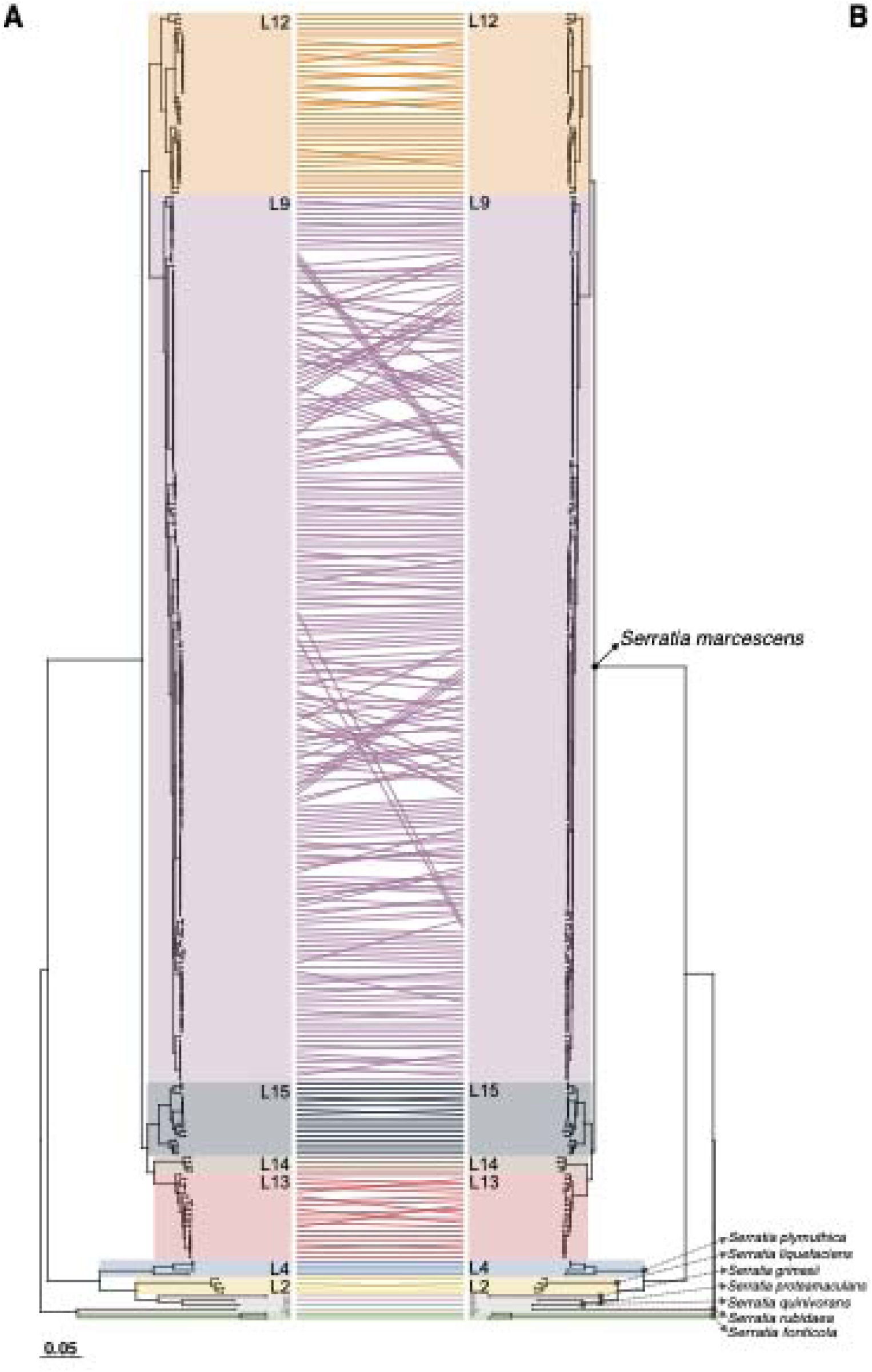
*PathogenSurveillance* core genome phylogeny reproduces a previously published phylogeny fully resolving species and lineages. The tanglegram shows congruence between (**A**) a maximum likelihood core gene phylogeny generated by *PathogenSurveillance*, and (**B**) a phylogeny reproduced from the literature. Matched samples are linked across trees by lines, with color-coding denoting lineage assignments. Major species and lineages (L#) are labeled as published. Scale bars represent the number of nucleotide substitutions per site. See narrative for reference citation.

To assess the accuracy of automated reference selection and taxonomic assignment by *PathogenSurveillance* we required a read dataset derived from samples with predictable taxonomic identities. We also required that no corresponding assemblies be available in public databases to ensure the pipeline will not automatically select them as references. Therefore, we generated a new Oxford Nanopore dataset from bacteria cultured from the guts of honeybees, which has a low complexity microbiota of only five core genera and is thus expected to yield strains of known taxonomic identity (Motta and Moran 2024). DNA sequenced from several colonies were preliminarily analyzed using *Kraken 2* and showed that most colonies were mixtures of strains (Wood et al. 2019). We selected two that were largely pure and, predicted to be from *Gilliamella apicola* strains by *Kraken 2*. The reads corresponding to *G. apicola* were extracted from these two samples and used to evaluate the efficacy of rules implemented for automatic reference selection (Supplementary Figure 2).

*PathogenSurveillance* automatically downloaded metadata for all possible reference assembles within the Orbales family, based on initial analysis by *Sendsketch*, and prioritized the candidate list according to a set of sorting criteria (listed in order of priority as columns in supplementary figure 2; (Bushnell 2014)). *PathogenSurveillance* then downloaded multiple references for each family, genus, and species for use as contextual references and for mapping. Importantly, the pipeline selected references for each of the six recognized *Gilliamella* species, Candidatus *Schmidhempelia*, and five other genera within the Orbaceae family. For *G. apicola, PathogenSurveillance* downloaded reference sequences from three distinct *G. apicola* strains, while avoiding those that lack key discriminating information, to maximize species-level diversity. Additionally, because *PathogenSurveillance* uses ANI as the key criterion to select mapping references, it selected the more similar *Gilliamella apicola* SCGC AB-598-B02 (GCF_000725165.1) over *Gilliamella apicola* (GCF_000599985.1), which was initially identified as the best representative of the species based on metadata (Supplementary Figure 2).

We also simulated sequence divergence of *G. apicola* samples to evaluate the robustness of reference selection. We simulated Illumina reads from the Oxford Nanopore-based assemblies and introduced up to 10% sequence divergence using the ART read simulator (Supplementary Figure 3)(Huang et al. 2012). Predictably, *PathogenSurveillance,* using simulated Illumina reads with no sequence divergence, selected the same references as the original dataset. Each increment of simulated sequence divergence, with the exception of 2% divergence, resulted in the selection of the same reference sequence. In the one exception, *PathogenSurveillance* selected *Gilliamella* sp. wkB7 (GCF_001693435.1), which is closely related and had highly similar ANI values. Therefore, the pipeline robustly selected references for phylogenetic inference and read mapping.

Lastly, we used publicly available genome sequences to generate prokaryotic and eukaryotic datasets to evaluate the impact of sample and genome sizes on runtime and RAM usage. One dataset was derived from a study of *Klebsiella pneumoniae*, which has an average genome size of 5.7 Mb (David et al. 2020)(Supplementary Table 3). To evaluate the effect of sample size, we randomly subsampled the dataset to form sets of 1, 3, 5, 10, 25, 50, 75, 100, 150, and 200 samples. The pipeline was executed with caching disabled to analyze three replicates of each of the sets. As expected, runtime showed a linear increase from 0.4 to 11.7 hours as the number of samples increased from 1 to 200 samples (Figure 4A). RAM also increased linearly from 4.3 GB for 1 sample to 12.2 GB for 200 samples (Figure 4B).

**Figure 4.**
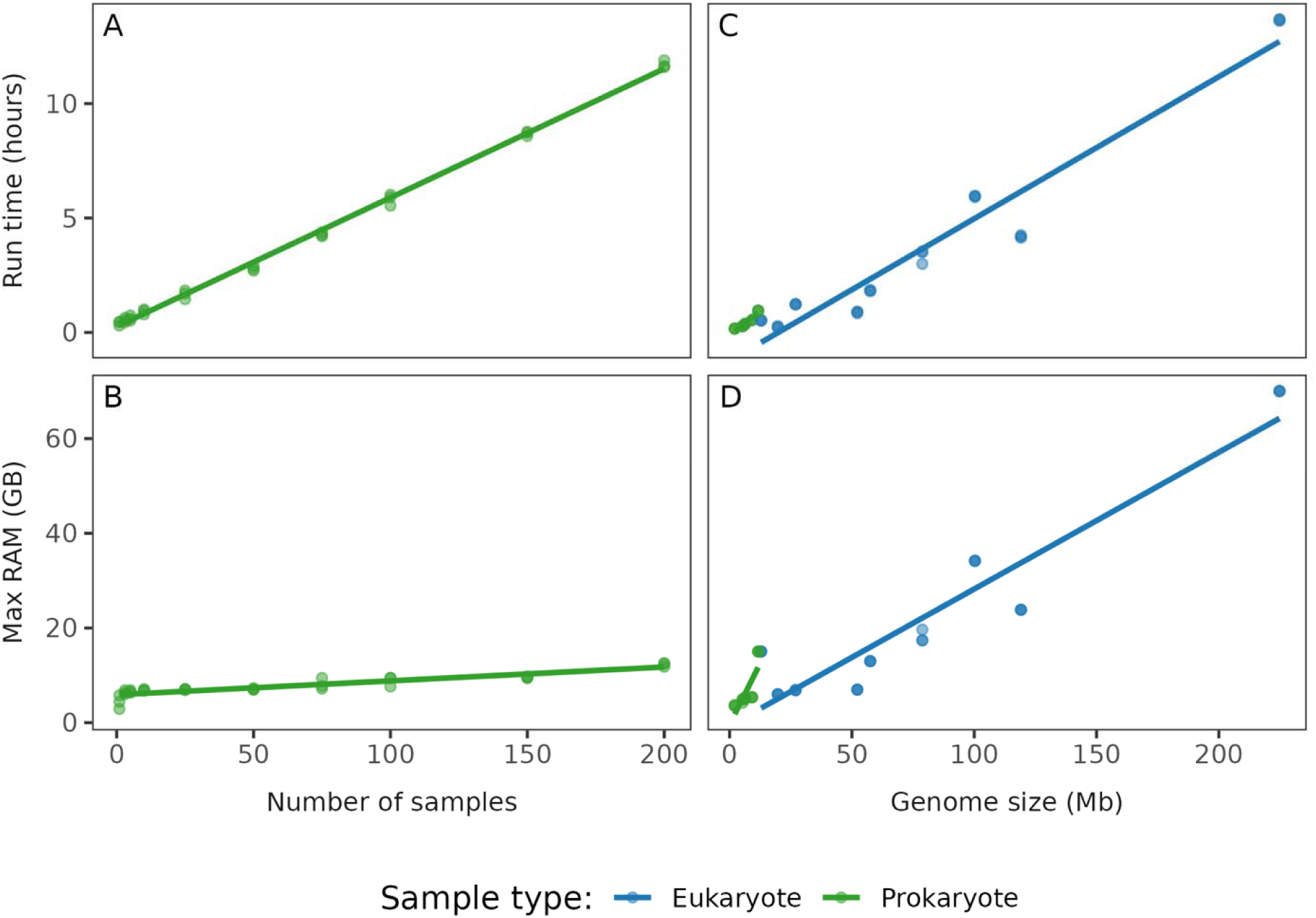
*PathogenSurveillance* demonstrates linear and scalable performance across a wide range of sample and genome sizes. Two test datasets were used to evaluate how the RAM and runtime of the pipeline scaled with number of samples and genome size. (**A**) and (**B**) show results for datasets increasing in sample sizes, while (**C**) and (**D**) show results for datasets with increasing genome sizes. Runtime (in clock hours) is plotted in (**A**) and (**C**), and maximum random-access memory (RAM) usage (in gigabytes (GB)) is plotted in (**B**) and (**D**). For genome size tests, each data set consisted of three samples of a given species and eukaryotes and prokaryotes are summarized separately since they are analyzed with different methods. Each dataset was analyzed in triplicate by *PathogenSurveillance*.

We also created a dataset of taxonomically disparate organisms with genome sizes that vary from 2.1 Mb to 224.8 Mb (David et al. 2020)(Supplementary Table 4). Each dataset was composed of three samples of a single species and was run in triplicate. Each consists of Illumina paired-end reads, which were subset to 25x coverage for prokaryotes and 50x for eukaryotes. The results for prokaryote and eukaryote genome sequences are displayed in different colors because they are processed by different workflow branches (Figure 1).

Prokaryote data, regardless of genome size, were each completed in less than one hour of automated analyses (Figure 4C) with RAM usage less than 15.0Gb (Figure 4D). In contrast, the runtime for eukaryotic genome sequences had a substantial range that scaled linearly to genome size. The largest genome took 13.6 hours of runtime. RAM usage ranged between 6.0 GB and 70.0 GB for the eukaryotic dataset.

All tests were run on a System 76 Thelio Major Linux Desktop with a Ryzen 9 7950x 16-core processor.

## Discussion

Pests and pathogens can emerge and rapidly adapt, often evading detection by traditional biosurveillance methods (Korber et al. 2020). In response to these challenges, we developed *PathogenSurveillance,* a user-friendly computational workflow for analyzing WGS data, with features specific for pest and pathogen detection. *PathogenSurveillance* provides a scalable computational workflow that is highly portable for implementation on any Linux cloud or workstation. Our implementation follows the FAIR principles. Importantly, the pipeline works for samples belonging to either known or unknown taxa and can resolve clonality or novel variants if WGS data is available for the clades in question.

A key design principle of *PathogenSurveillance* is its commitment to lowering the barrier to entry and accelerating the analyses of WGS data. Population-level analysis requires domain specific knowledge in molecular evolution and bioinformatics, particularly for selecting appropriate references, installing software and dependencies, and constructing robust phylogenies. At its core, *PathogenSurveillance* is a generalizable population genomics pipeline, featuring a rules-based system that automates reference selection. Leveraging the Nextflow workflow management system, this design empowers users from diverse backgrounds, including those without formal training in molecular evolution and bioinformatics, to effectively analyze WGS data. Additionally, the organization of results reflects a deliberate design intended to support further in-depth investigations, both through manual and automated approaches. Collectively, *PathogenSurveillance* provides a streamlined and accessible framework for automating initial population genomic analysis, supporting both applied and fundamental research applications.

There are constraints to *PathogenSurveillance* that should be considered. First, the pipeline does not support analyses of viruses, as they would require their own workflow branch to parse metagenomic data and employ specialized approaches for taxonomic identification (Rangel-Pineros et al. 2023). Second, the accuracy in which the pipeline predicts taxonomic identities is constrained by the contents of databases used and the correctness of the taxonomic names they contain. Third, *PathogenSurveillance* was designed for WGS data derived from cultured organisms, which may limit applicability to metagenomic or environmental samples. Fourth, *PathogenSurveillance* employs default parameters across all steps, which may require appropriate adjustments for non-standard cases or refining reference selection. Finally, the pipeline can be computationally demanding. While the Nextflow framework allows efficient memory allocation per task, certain processes and samples may require substantial memory.

Moreover, the pipeline generates numerous intermediate files, requiring storage capacity that is approximately five times greater than the size of input files. Nonetheless, there are ways to apply *PathogenSurveillance* that mitigate these constraints. For example, we have identified contaminating reads in some sample datasets, which is analogous to a low complexity metagenome dataset. When left unfiltered, the pipeline successfully resolved the taxonomic identity of the primary organism, suggesting potential use in metagenome analyses. Regarding use of default parameters, caching gives users the flexibility of running the pipeline with a preliminary assessment and rerunning it with refined parameters or references without repeating the entire analysis. This not only saves considerable time but also supports reproducibility through automated, consistent execution. To mitigate storage requirements, users can clear the cache after analysis. However, for taxa that are repeatedly analyzed, retaining cached results may be advantageous, as it offsets rerun times.

In summary, *PathogenSurveillance* automates the analyses of WGS data with a single command. It enhances traditional diagnostic approaches by improving analytical resolution and enriching the data available for more robust interpretation. It can also aid in identifying novel or emerging pathogens as well as place variants within taxa. These features underscore the potential use of *PathogenSurveillance* in global pathogen and pest biosurveillance (Carvajal-Yepes et al. 2019; Weisberg et al. 2021).

## Data and Source Code Availability

*PathogenSurveillance* is free and open source under the MIT license. The source code is hosted on Github (https://github.com/nf-core/PathogenSurveillance) and documentation is available on nf-core (https://nf-co.re/PathogenSurveillance/). All data used in the validation of the pipeline presented here are available in the test data repository on GitHub (https://github.com/nf-core/test-datasets/tree/PathogenSurveillance). The scripts and input data used to create figures in this manuscript are also available on GitHub (https://github.com/grunwaldlab/PathogenSurveillance_publications). Test datasets consisting of sequences that are automatically downloaded from NCBI are available as profiles and can easily be run as documented at https://github.com/nf-core/test-datasets/tree/PathogenSurveillance. Supplementary tables and figures are found at https://github.com/grunwaldlab/pathogensurveillance_publications/tree/main/publication/supplementary_data.

## Acknowledgements

We thank members of the Chang and Grünwald laboratories for their valuable input and support. We also thank Melodie Putnam for her thoughtful advice on the design principles of *PathogenSurveillance,* Dr. Maude David for providing bacteria of the honeybee microbiota, and Charles Paulsen for extracting DNA and making libraries. We also thank the nf-core community for developing the infrastructure and resources for Nextflow pipelines, in particular Harshil Patel, Phil Ewels, Matthias Hörtenhuber, Mahesh Binzer-Panchal, and Maxime Garcia, for their assistance in reviewing and releasing the pipeline. Lastly, we thank the Center for Quantitative Life Sciences (CQLS) and Department of Botany and Plant Pathology (BPP) at Oregon State University and SCINet at the United States Department of Agriculture – Agricultural Research Service (USDA-ARS) for supporting high-performance computing infrastructures.

## Author Contributions

ZSLF designed and implemented nf-core/*PathogenSurveillance*. MAS, CPR, LKB, FIB, AJW, and HP contributed scripts and modules, and/or enhanced or tested pipeline reliability, usability and performance, and TRS cultured bacteria from honeybees. ZSLF, MAS, CPR, JHC, and NJG wrote the manuscript, with input from all authors. NJG and JHC led the conceptualization and planning of nf-core/*PathogenSurveillance*, supervised its development, and secured funding.

## Supplementary data

**Supplementary Figure 1.**
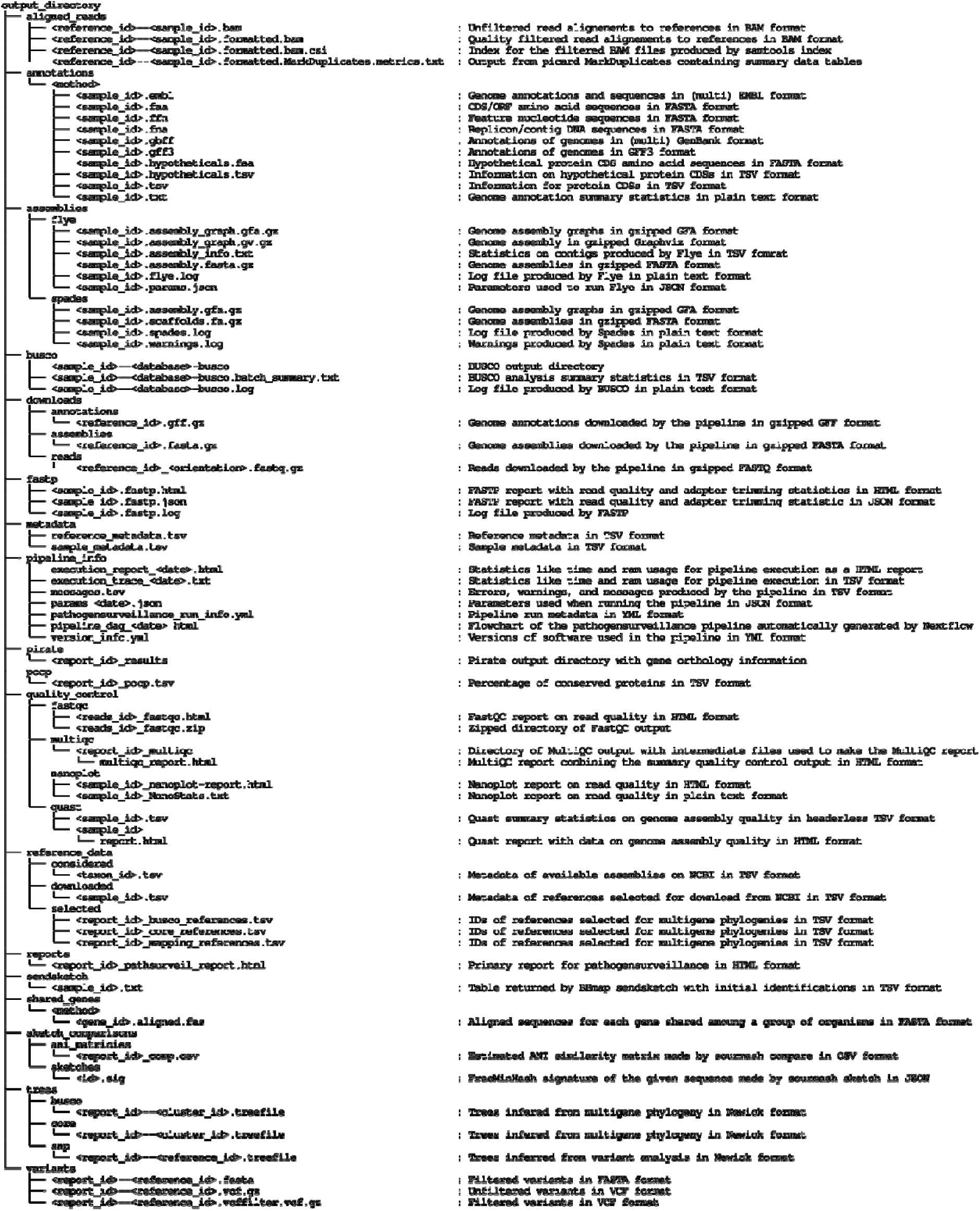
*PathogenSurveillance* organizes and names output files for further analyses. The left side displays the hierarchical structure of output files. Angle brackets (<>) show run specific identifiers for references, samples, and reports. The right side provides brief descriptions of each file type, highlighting their applications in downstream analyses.

**Supplementary Figure 2.**
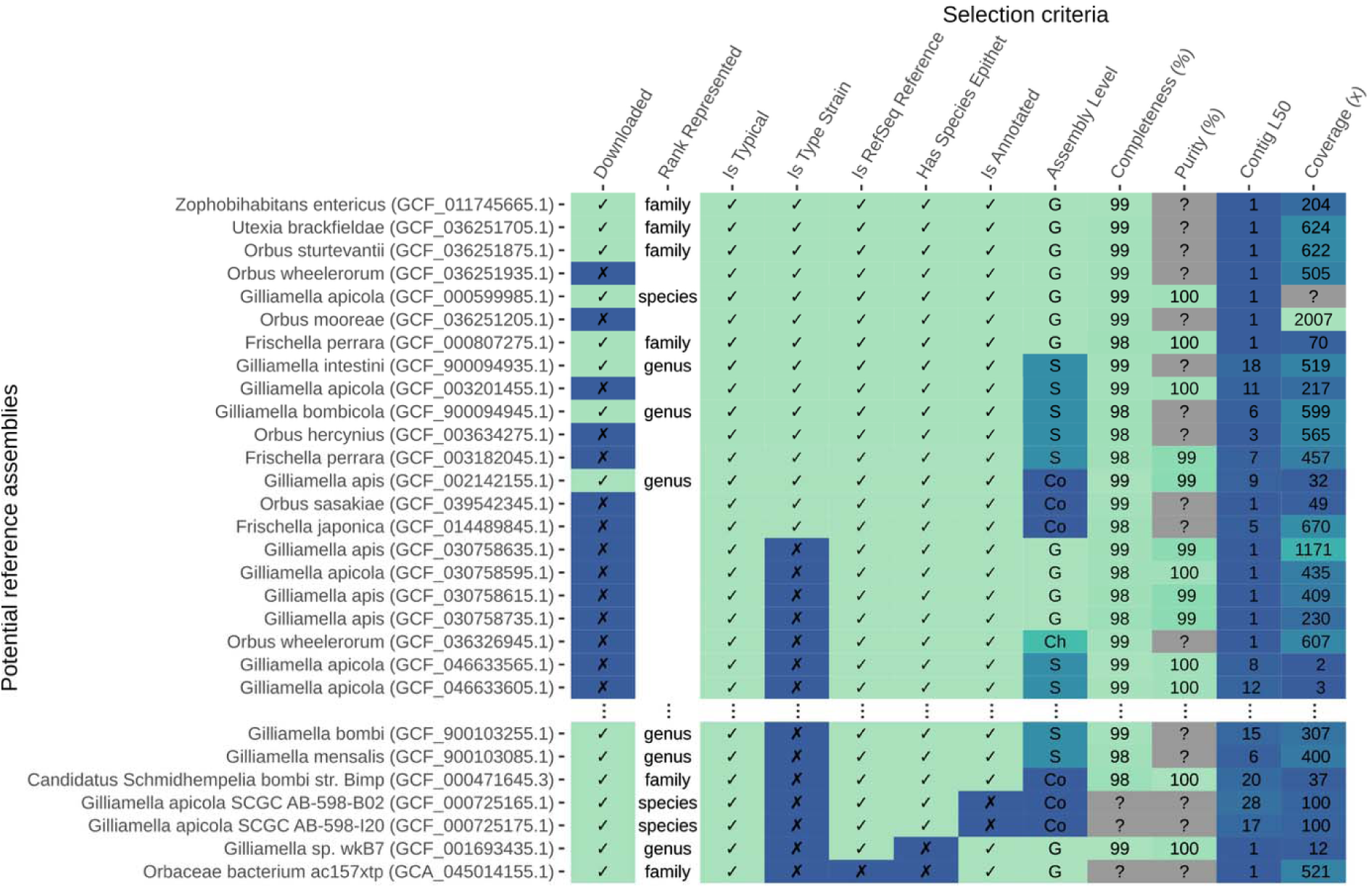
*PathogenSurveillance* automates downloading of references based on taxa found in the initial identification. Representatives of taxa predicted by sendsketch are chosen based on available metadata relevant to assembly quality and downloaded. This shows the criteria used to choose references for two samples of *Gilliamella apicola*. Representatives of the species, genus, and family are chosen, in case the species-level identification is incorrect. Potential references are represented by rows and sorted from best to worst based on the selection criteria represented by columns. Selection criteria columns are sorted from most important to least important.

**Supplementary Figure 3.**
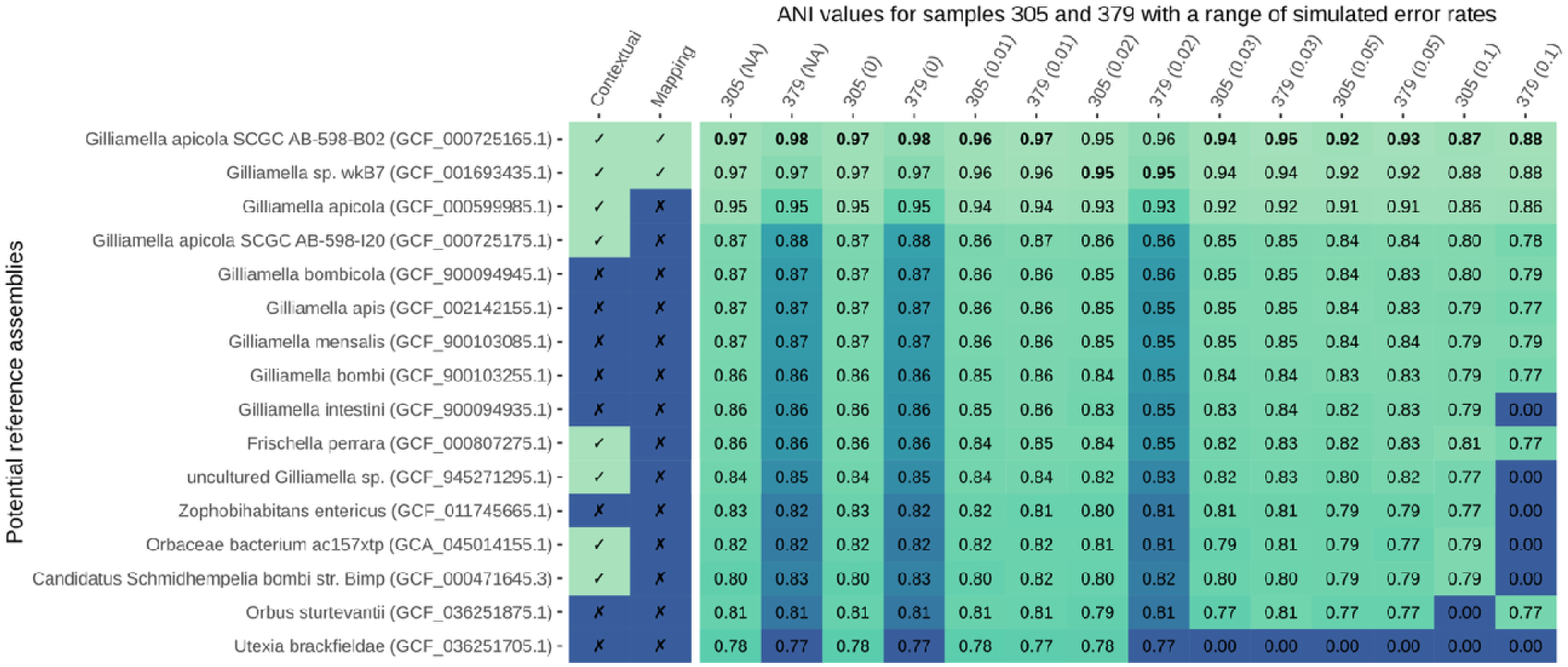
*PathogenSurveillance* automates selection of contextual and mapping references from those downloaded. Nanopore reads of two samples of *Gilliamella apicola* (305 and 379) were used as test data. Short reads were simulated from the nanopore reads with a range of error rates (indicated in the parentheses in column names starting third column). *Pathogensurveillance* was run on each pair of samples from which references were selected for multigene phylogenies (contextual) and variant calling (mapping). ANI values estimated by sourmash are shown in cells and bolded ANI values represent which mapping reference was chosen for each pair of samples.

## References

Bushnell B. 2014. BBMap: a fast, accurate, splice-aware aligner. Available from: https://www.osti.gov/biblio/1241166

Carvajal-Yepes M, Cardwell K, Nelson A, Garrett KA, Giovani B, Saunders DGO, Kamoun S, Legg JP, Verdier V, Lessel J, et al. 2019. A global surveillance system for crop diseases. Science 364:1237–1239.

Chen S, Zhou Y, Chen Y, Gu J. 2018. fastp: an ultra-fast all-in-one FASTQ preprocessor. Bioinformatics 34:i884–i890.

David S, Cohen V, Reuter S, Sheppard AE, Giani T, Parkhill J, the European Survey of Carbapenemase-Producing Enterobacteriaceae (EuSCAPE) Working Group, the ESCMID Study Group for Epidemiological Markers (ESGEM), Rossolini GM, Feil EJ, et al. 2020. Integrated chromosomal and plasmid sequence analyses reveal diverse modes of carbapenemase gene spread among *Klebsiella pneumoniae*. Proc. Natl. Acad. Sci. U.S.A. 117:25043–25054.

De Coster W, Rademakers R. 2023. NanoPack2: population-scale evaluation of long-read sequencing data. Alkan C, editor. Bioinformatics 39:btad311.

De Visser C, Johansson LF, Kulkarni P, Mei H, Neerincx P, Joeri Van Der Velde K, Horvatovich P, Van Gool AJ, Swertz MA, Hoen PAC ‘T, et al. 2023. Ten quick tips for building FAIR workflows. Palagi PM, editor. PLoS Comput Biol 19:e1011369.

Di Tommaso P, Chatzou M, Floden EW, Barja PP, Palumbo E, Notredame C. 2017. Nextflow enables reproducible computational workflows. Nat Biotechnol 35:316–319.

Djordjevic SP, Jarocki VM, Seemann T, Cummins ML, Watt AE, Drigo B, Wyrsch ER, Reid CJ, Donner E, Howden BP. 2024. Genomic surveillance for antimicrobial resistance — a One Health perspective. Nat Rev Genet 25:142–157.

Eggertsson HP, Kristmundsdottir S, Beyter D, Jonsson H, Skuladottir A, Hardarson MT, Gudbjartsson DF, Stefansson K, Halldorsson BV, Melsted P. 2019. GraphTyper2 enables population-scale genotyping of structural variation using pangenome graphs. Nat Commun 10:5402.

Ewels P, Magnusson M, Lundin S, Käller M. 2016. MultiQC: summarize analysis results for multiple tools and samples in a single report. Bioinformatics 32:3047–3048.

Fisher MC, Henk DanielA, Briggs CJ, Brownstein JS, Madoff LC, McCraw SL, Gurr SJ. 2012. Emerging fungal threats to animal, plant and ecosystem health. Nature 484:186–194.

Gardy JL, Loman NJ. 2018. Towards a genomics-informed, real-time, global pathogen surveillance system. Nat Rev Genet 19:9–20.

Grünwald NJ, Goss EM. 2011. Evolution and population genetics of exotic and re-emerging pathogens: Novel tools and approaches. VanAlfen, NK and Bruening, G and Leach, JE, editor. Annual Review of Phytopathology 49:249–267.

Gurevich A, Saveliev V, Vyahhi N, Tesler G. 2013. QUAST: quality assessment tool for genome assemblies. Bioinformatics 29:1072–1075.

Hamelin RC, Bilodeau GJ, Heinzelmann R, Hrywkiw K, Capron A, Dort E, Dale AL, Giroux E, Kus S, Carleson NC, et al. 2022. Genomic biosurveillance detects a sexual hybrid in the sudden oak death pathogen. Commun Biol 5:477.

Huang W, Li L, Myers JR, Marth GT. 2012. ART: a next-generation sequencing read simulator. Bioinformatics 28:593–594.

Hulme PE. 2009. Trade, transport and trouble: managing invasive species pathways in an era of globalization. Journal of Applied Ecology 46:10–18.

Iles LC, Fulladolsa AC, Smart A, Bonkowski J, Creswell T, Harmon CL, Hammerschmidt R, Hirch RR, Rodriguez Salamanca L. 2021. Everything is faster: How do land-grant university–based plant diagnostic laboratories keep up with a rapidly changing world? Annu. Rev. Phytopathol. 59:333–349.

Irber L, Pierce-Ward NT, Abuelanin M, Alexander H, Anant A, Barve K, Baumler C, Botvinnik O, Brooks P, Dsouza D, et al. 2024. sourmash v4: A multitool to quickly search, compare,and analyze genomic and metagenomic data sets. JOSS 9:6830.

Irber L, Pierce-Ward NT, Brown CT. 2022. Sourmash branchwater enables lightweight petabyte-scale sequence search. bioRxiv:2022.11.02.514947.

Iruegas-Bocardo F, Weisberg AJ, Riutta ER, Kilday K, Bonkowski JC, Creswell T, Daughtrey ML, Rane K, Grünwald NJ, Chang JH, et al. 2023. Whole genome sequencing-based tracing of a 2022 introduction and outbreak of *Xanthomonas hortorum* pv. *pelargonii*. Phytopathology® 113:975–984.

Kamvar ZN, Tabima JF, Grünwald NJ. 2014. *Poppr*: an R package for genetic analysis of populations with clonal, partially clonal, and/or sexual reproduction. PeerJ 2:e281.

Konstantinidis KT, Tiedje JM. 2005. Genomic insights that advance the species definition for prokaryotes. Proc. Natl. Acad. Sci. U.S.A. 102:2567–2572.

Korber B, Fischer WM, Gnanakaran S, Yoon H, Theiler J, Abfalterer W, Hengartner N, Giorgi EE, Bhattacharya T, Foley B, et al. 2020. Tracking Changes in SARS-CoV-2 Spike: Evidence that D614G Increases Infectivity of the COVID-19 virus. Cell 182:812–827.e19.

Liebhold AM, Brockerhoff EG, Garrett LJ, Parke JL, Britton KO. 2012. Live plant imports: the major pathway for forest insect and pathogen invasions of the US. Frontiers in Ecology and the Environment 10:135–143.

Manni M, Berkeley MR, Seppey M, Simão FA, Zdobnov EM. 2021. BUSCO Update: Novel and Streamlined Workflows along with Broader and Deeper Phylogenetic Coverage for Scoring of Eukaryotic, Prokaryotic, and Viral Genomes. Kelley J, editor. Molecular Biology and Evolution 38:4647–4654.

Meyerson LA, Mooney HA. 2007. Invasive alien species in an era of globalization. Frontiers in Ecology and the Environment 5:199–208.

Motta EVS, Moran NA. 2024. The honeybee microbiota and its impact on health and disease. Nat Rev Microbiol 22:122–137.

Oakeson KF, Wagner JM, Mendenhall M, Rohrwasser A, Atkinson-Dunn R. 2017. Bioinformatic analyses of whole-genome sequence data in a public health laboratory. Emerg. Infect. Dis. 23:1441–1445.

Ondov BD, Treangen TJ, Melsted P, Mallonee AB, Bergman NH, Koren S, Phillippy AM. 2016. Mash: fast genome and metagenome distance estimation using MinHash. Genome Biol 17:132.

Petit RA, Read TD. 2020. Bactopia: a Flexible Pipeline for Complete Analysis of Bacterial Genomes. Segata N, editor. mSystems 5:e00190–20.

Pierce NT, Irber L, Reiter T, Brooks P, Brown CT. 2019. Large-scale sequence comparisons with sourmash. F1000Res 8:1006.

Qin Q-L, Xie B-B, Zhang X-Y, Chen X-L, Zhou B-C, Zhou J, Oren A, Zhang Y-Z. 2014. A Proposed Genus Boundary for the Prokaryotes Based on genomic insights. J Bacteriol 196:2210–2215.

Rangel-Pineros G, Almeida A, Beracochea M, Sakharova E, Marz M, Reyes Muñoz A, Hölzer M, Finn RD. 2023. VIRify: An integrated detection, annotation and taxonomic classification pipeline using virus-specific protein profile hidden Markov models.Ouzounis CA, editor. PLoS Comput Biol 19:e1011422.

Tian L, Huang C, Mazloom R, Heath LS, Vinatzer BA. 2020. LINbase: a web server for genome-based identification of prokaryotes as members of crowdsourced taxa. Nucleic Acids Research 48:W529–W537.

Weisberg AJ, Grünwald NJ, Savory EA, Putnam ML, Chang JH. 2021. Genomic approaches to plant-pathogen epidemiology and diagnostics. Annu. Rev. Phytopathol. 59:311–332.

Williams DJ, Grimont PAD, Cazares A, Grimont F, Ageron E, Pettigrew KA, Cazares D, Njamkepo E, Weill F-X, Heinz E, et al. 2022. The genus *Serratia* revisited by genomics. Nat Commun 13:5195.

Wood DE, Lu J, Langmead B. 2019. Improved metagenomic analysis with Kraken 2. Genome Biol 20:257.

